# The Arabidopsis Auxin Receptor F-box proteins AFB4 and AFB5 are Required for Response to the Synthetic Auxin Picloram

**DOI:** 10.1101/034652

**Authors:** Michael J. Prigge, Kathleen Greenham, Yi Zhang, Aaron Santner, Cristina Castillejo, Ronan C. O’Malley, Joseph R. Ecker, Mark Estelle

## Abstract

The plant hormone auxin is perceived by a family of F-box proteins called the TIR1/AFBs. Phylogenetic studies reveal that these proteins fall into four clades in flowering plants called TIR1, AFB2, AFB4, and AFB6 (Parry *et al*. 2009). Genetic studies indicate that members of the TIR1 and AFB2 groups act as positive regulators of auxin signaling by promoting the degradation of the Aux/IAA transcriptional repressors (Dharmasiri *et al*. 2005; Parry *et al*. 2009). In this report, we demonstrate that both AFB4 and AFB5 also function as auxin receptors based on *in vitro* assays. We also provide genetic evidence that both AFB4 and AFB5 are targets of the picloram family of auxinic herbicides. In contrast to previous studies we find that null *afb4* alleles do not exhibit obvious defects in seedling morphology or auxin hypersensitivity. We conclude that AFB4 and AFB5 act in a similar fashion to other members of the family but exhibit a distinct auxin specificity.

## Introduction

The plant hormone auxin is a small indolic molecule with an important role in virtually every aspect of plant growth and development from embryogenesis to senescence (Woodward and Bartel 2005). Auxin regulates transcription by promoting the degradation of a family of transcriptional repressors called the Aux/IAA proteins (Hagen 2015; Salehin *et al*. 2015). These proteins repress transcription by binding to transcription factors called AUXIN RESPONSE FACTORs (ARFs), and recruiting the co-repressor protein TOPLESS to the chromatin. In the presence of auxin, the AUXIN/INDOLE-3-ACETIC ACID (Aux/IAA) proteins are degraded through the action of a ubiquitin protein ligase (E3) called SCF^TIR1^. This results in activation of complex transcriptional networks that lead to context-dependent changes in cell growth and behavior.

The SCFs are a subgroup of a large family of E3 ligases called Cullin Ring Ligases (CRL) conserved in all eukaryotes (Pickart 2001; Petroski and Deshaies 2005). SCFs consist of CULLIN1, S-phase kinase associated protein 1 (SKP1,ASK in plants), the RING-BOX1 (RBX1) protein, and one of a family of substrate adaptor proteins called F-box proteins (Pickart 2001; Petroski and Deshaies 2005). The F-box protein recruits substrates to the SCF and promotes ubiquitination, typically resulting in degradation by the proteasome. Several years ago, we discovered that SCF^TIR1^ and the related SCF^AFBs^ function as auxin sensors (Dharmasiri *et al*. 2005; Kepinski and Leyser 2005; Tan *et al*. 2007). The TRANSPORT INHIBITOR RESPONSE1/AUXIN F-BOX (TIR1/AFB) proteins consist primarily of 18 Leucine Rich Repeats (LRRs). Auxin binds directly to the LRR region of TIR1, but rather than causing a conformational change, typical for most hormone receptors, auxin promotes the interaction between SCF^TIR1^ and the Aux/IAA substrate.

There are six members of the TIR1/AFB group of F-box proteins in Arabidopsis. TIR1 and AFB1 through AFB3, as well as AFB5 have been shown to function as auxin receptors (Dharmasiri *et al*. 2005; Calderon Villalobos *et al*. 2012). The loss of a single member of *TIR1* through *AFB3* has a slight effect on auxin response and plant growth, but higher order combinations of these genes have a much more severe phenotype (Dharmasiri *et al*. 2005). Of these four proteins TIR1 and AFB2 appear to have major roles in seedling development, while AFB3 has a less significant role. The loss of AFB1 has a very minor effect in the seedling (Dharmasiri *et al*. 2005). This appears to be due to the fact that AFB1 does not assemble into an SCF complex efficiently (Yu *et al*. 2015). In this study we focus on the *AFB4* and *AFB5* genes. We describe the characterization of two new *AFB4* mutants called *afb4-8* and *afb4-9*. Both of these mutations appear to be null alleles, but neither has an obvious effect on growth of the seedling. We confirm that both AFB4 and AFB5 function as auxin receptors. In addition we show that the *afb4* and *afbs* mutants are resistant to the synthetic auxin picloram indicating that these two proteins are selective for picloram.

## Experimental Procedures

### Plant material and growth conditions and treatments

*Arabidopsis thaliana* mutants and transgenic lines used in this study were all in the Columbia (Col-0) ecotype. The Salk T-DNA insertion lines *afb4-8* (Salk_201329) and *afb4-9* (Salk_083223) were identified in the Salk-seq data (http://signal.salk.edu/cgi-bin/tdnaexpress). The *afb4-9* line originally contained four additional T-DNA insertions. A previously described *At5g27570/cdc20.5* insertion (Kevei *et al*. 2011) and an insertion in the *At1g11340* gene were removed by backcrossing, but two intergenic insertions near genes *At3g09720* and *At4g22160* remained present in the *afb4-9* and *afb4-9 afb5-5* lines used in this study. The *afb5-5* (Salk_110643) was obtained from the Arabidopsis Biological Resource Center at Ohio State University. The plant-T-DNA junction sequences were determined for each insertion. The *afb4-8* insertion is associated with a 20 bp deletion, while those of *afb4-9* and *afb5-5* are 10 bp and 32 bp, respectively. Seeds were surface sterilized either by vapor-phase sterilization (Clough and Bent 1998) or by treating for 2 min in 70% (v/v) ethanol followed by 10 min in 30% commercial bleach. Seeds were plated on medium containing 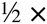 MS media, 1% sucrose, 0.8% agar and stratified for 2-4 days at 4°C.

### Growth Assays

All root assays were completed under long-day photoperiods (16:8) and hypocotyl assays were performed under short-day photoperiods (8:16). For auxin-inhibited root growth assays, 5-day-old seedlings were transferred onto fresh MS media ± auxin for 3 additional days after which root length was measured. Hypocotyl assays were performed similarly except the seedlings were transferred at day 4 for a 2-day treatment.

### Generation of Transgenic Lines

The *TIR1-Myc* line was described previously (Gray *et al*. 1999). The AFB4- and *AFB5-4×Myc* lines were generated using a 2-kb 5’ upstream region of the *AFB5* gene with the *AFB4* and *AFB5* cDNA in binary vector pGW 16. The *AFB5* promoter was used for expressing AFB4 due to the low activity of the *AFB4* promoter. The *AFB5-mCitrine* construct contained the entire genomic region between adjacent genes, from 1267 bp upstream of the start codon to 1139 bp downstream from the stop codon in the pMP535 binary vector (Prigge *et al*. 2005) The stop codon was mutated to a *Nhe*I site in order to insert a 27 bp linker and the mCitrine coding region. Each construct was transformed into the *afb5-5* mutant background.

### Protein Expression and Pulldown experiments

For pulldown assays, GST-IAA3 and GST-IAA7 were recombinantly expressed in *E. coli* strains BL21(DE3) (Figures 1B, 4C, and 4D) or BL21-AI (Figure 5) and purified using Glutathione-Agarose (Sigma-Aldrich, G4510). For *in vivo* pulldown experiments, seedlings expressing Myc-tagged AFB4, AFB5 and TIR1 were grown for 8 days in liquid MS medium. TIR1-Myc expression was induced by treatment with 30μM Dex for 24h. The ASK1-antibody was generated as previously described (Gray *et al*. 1999). For the various auxin comparisons (Figure 4B) seedlings were incubated for 2h in 50μM of the compounds or an equivalent volume of DMSO prior to harvest. For all other *in vivo* pulldown experiments samples were incubated with auxin for 45min following harvest. Tissue was harvested by grinding to a powder in liquid nitrogen and vortexed vigorously in extraction buffer (50mM Tris pH7.5, 150mM NaCl, 10% glycerol, 0.1% NP-40, complete protease inhibitor (Roche), 50μM MG-132). Cellular debris was removed by centrifugation and total protein concentration was determined by Bradford assay. Each pulldown reaction included 1mg total protein extract and equal volumes of GST-IAA protein for each sample in a 500μl total volume. The pulldown reactions were incubated at 4°C for 45min with rocking and transferred to a Micro Bio-Spin Chromatography Column (Bio-Rad). Samples were washed 3 times in 1 ml extraction buffer without protease inhibitors or MG-132 in the presence or absence of auxin. Samples were eluted using reduced glutathione (Sigma) and separated on SDS-PAGE and stained with Ponceau (0.1% (w/v) Ponceau S in 5%(v/v) acetic acid) for loading control unless otherwise indicated. For Figure 4B, equivalent amounts were run on a separate SDS-PAGE gel and stained with Coomassie stain. AFB/TIR1-Myc proteins were detected by immunoblotting with anti-c-Myc-Peroxidase antibody (Roche). Proteins were visualized using the ECL Plus Western Blotting Detection System (Amersham).

**Figure 1.**
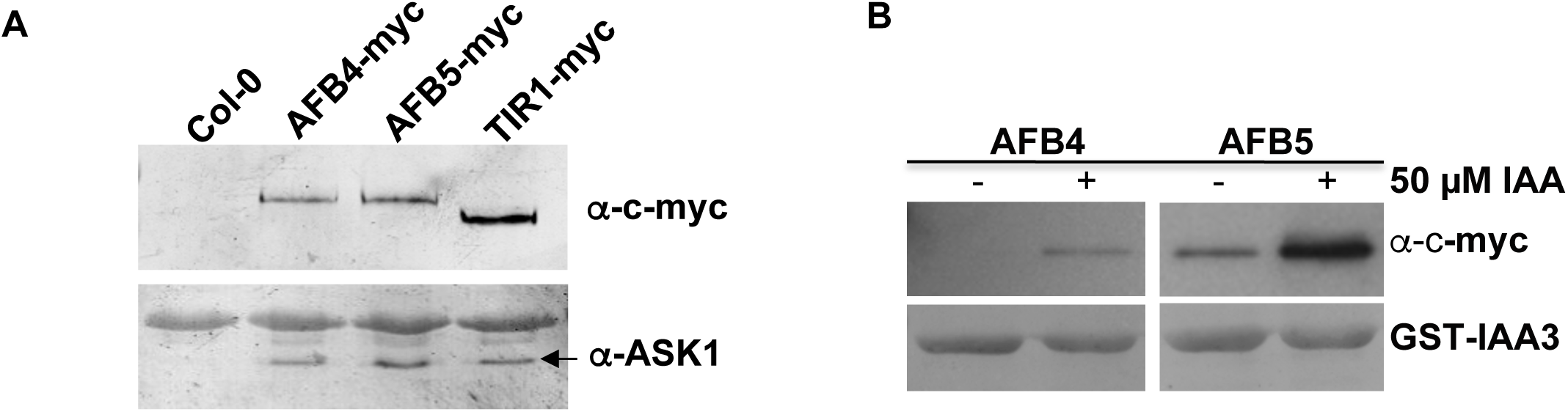
AFB4 and AFB5 interact with ASK1 and interact with the Aux/IAAs in an auxin dependent manor revealing their role as auxin receptors. (A-B) Pull-down experiments were carried out using crude plant extracts prepared from *[tir1-1] GVG> TIR1-Myc, [afb5-5] P_AFB5_:AFB5-Myc* and *[afb5-5] P_AFB5_:AFB4-Myc* seedlings and recombinant GST-IAA3. (A) TIR1-Myc, AFB4-Myc and AFB5-Myc were immunoprecipitated with the anti-Myc antibody coupled to agarose beads, and ASK1 was detected with an anti-ASKl antibody. (B) GST-IAA3 was immunoprecipitated with glutathione agarose beads, and AFB4-Myc and AFB5-Myc protein were detected with the anti-c-Myc-Peroxidase antibody. Pull-down reactions were incubated for 45min in the presence or absence of 5CμM IAA.

For the *in vitro* pulldown experiment, expression plasmids were made by adding the *AFB4* and *afb4^D215N^* cDNA sequences to a pTNT vector (Promega) with a Gateway:4×Myc cassette via Gateway recombination (Invitrogen). AFB4-4×Myc, afb4^D215N^-4×Myc and TIR1-Myc were produced from TNT T7 coupled wheat germ extract system (Promega, L4140). Comparable amount of AFB4-Myc, afb4^D215N^-Myc and TIR1-Myc were applied to each pull-down reaction as guided by western blot using anti-c-Myc-Peroxidase antibody (Roche, 11814150001). The pull-down assay was performed as described in Yu et al., 2013. TNT products and GST-IAA7 beads were incubated with or without the addition of 50μM IAA. The eluted products were detected and visualized as with the *in vivo* pulldowns.

### RNA extraction and quantitative PCR

Hypocotyl, cotyledon and root tissue frozen in liquid N_2_ and ground using a mortar and pestle was used for RNA purification using the Invitrogen PureLink RNA minikit. RNA from whole 10-day old seedlings (Figure 2C) was similarly ground and purified using RNeasy Plant Mini kit (Qiagen). RNA yield was quantified using the Thermo Scientific NanoDrop 2000. For quantitative RT-PCR, 1μg RNA, pre-treated with DNase using the DNA-free Kit (Ambion) according to manufacturer’s instructions, was used for generating cDNA with SuperScript IV (Figure 2) or SuperScript III (Figure 6A) Reverse Transcriptase (Invitrogen) and 20-mer oligo(dT) primers. Quantitative RT-PCR was performed using SyBR green and the primers listed in Table 1. Primer pairs were evaluated for specificity and efficiency using three serial dilutions of cDNA using the CFX96^TM^ Real-Time PCR Detection System (Biorad). Most data were normalized to the reference primer pair PP2AA3-S (Czechowski *et al*. 2005) according to the ΔΔCt method. Primer pairs AFB4-3 and AFB5-2 were normalized to the reference primer pair PP2AA3-L primer pair. All new primers were designed using QuantPrime (Arvidsson *et al*. 2008). Two biological replicates were performed, each replicate containing 95 to 100 mg whole seedlings (Figure 2C) or roughly 700 individual seedlings that were dissected into cotyledon, hypocotyl and root samples (Figure 6A).

**Figure 2.**
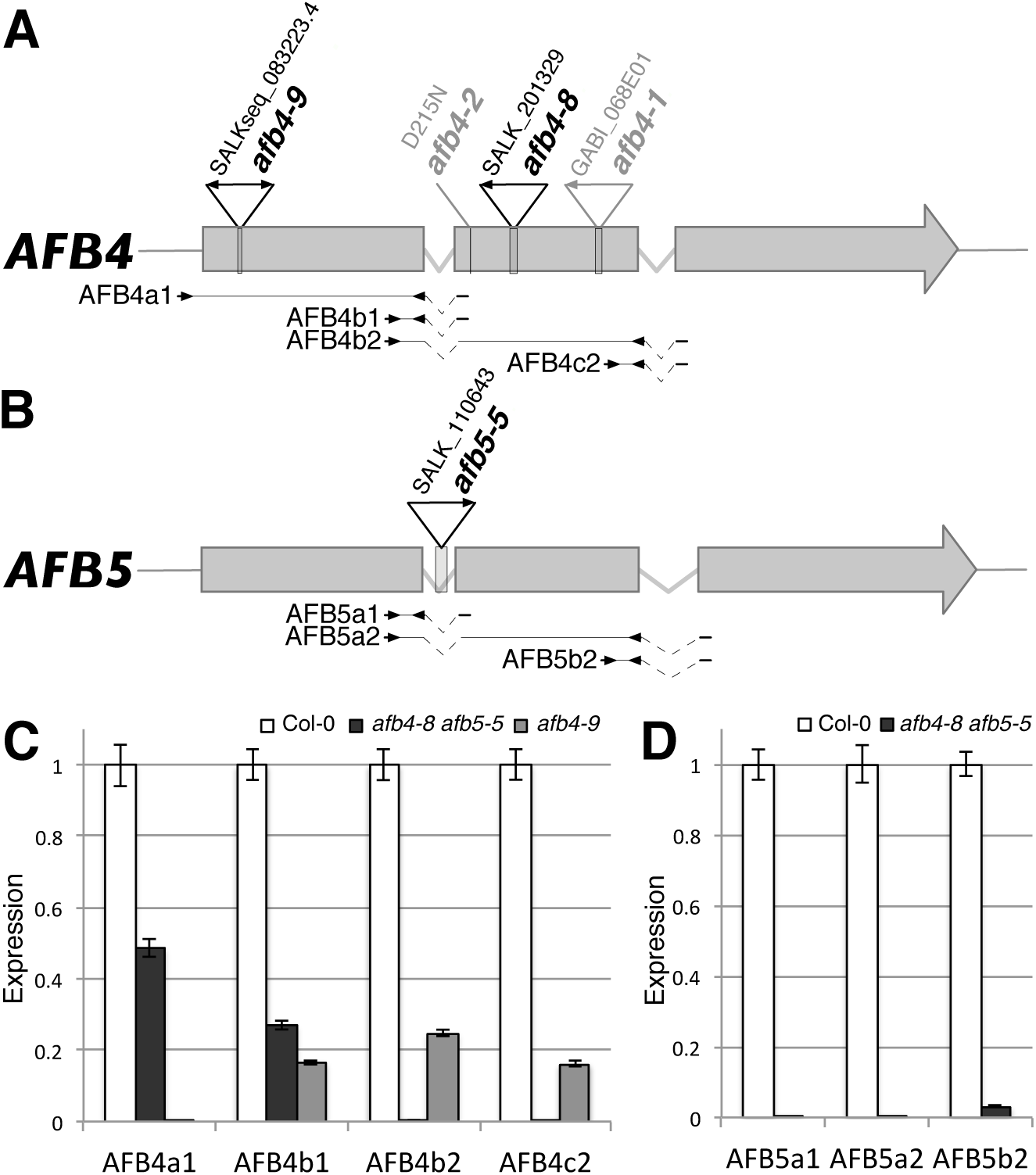
*afb4-8, afb4-9*, and *afb5-5* mutants do not produce full-length transcripts. (A-B) Diagrams of the *AFB4* and *AFB5* genes. The positions of mutant lesions are shown above the genes with arrowheads indicating T-DNA left border sequences. Below the gene diagrams are the primers pairs used for qRT-PCR. Kinked dashed lines indicate spliced introns. (C-D) qRT-PCR of *AFB4* and *AFB5* transcripts in WT and mutants grown under LD conditions. Results from each *AFB4* and *AFB5* primer pairs were normalized relative to those to the *PP2AA3* gene. AFB4a1, AFB4b2, and AFB5a2 were normalized to the longer PP2AA3-L amplicon (448 bp) while the rest used PP2AA3-S (59 bp). Error bars represent standard error.

**Table 1.**
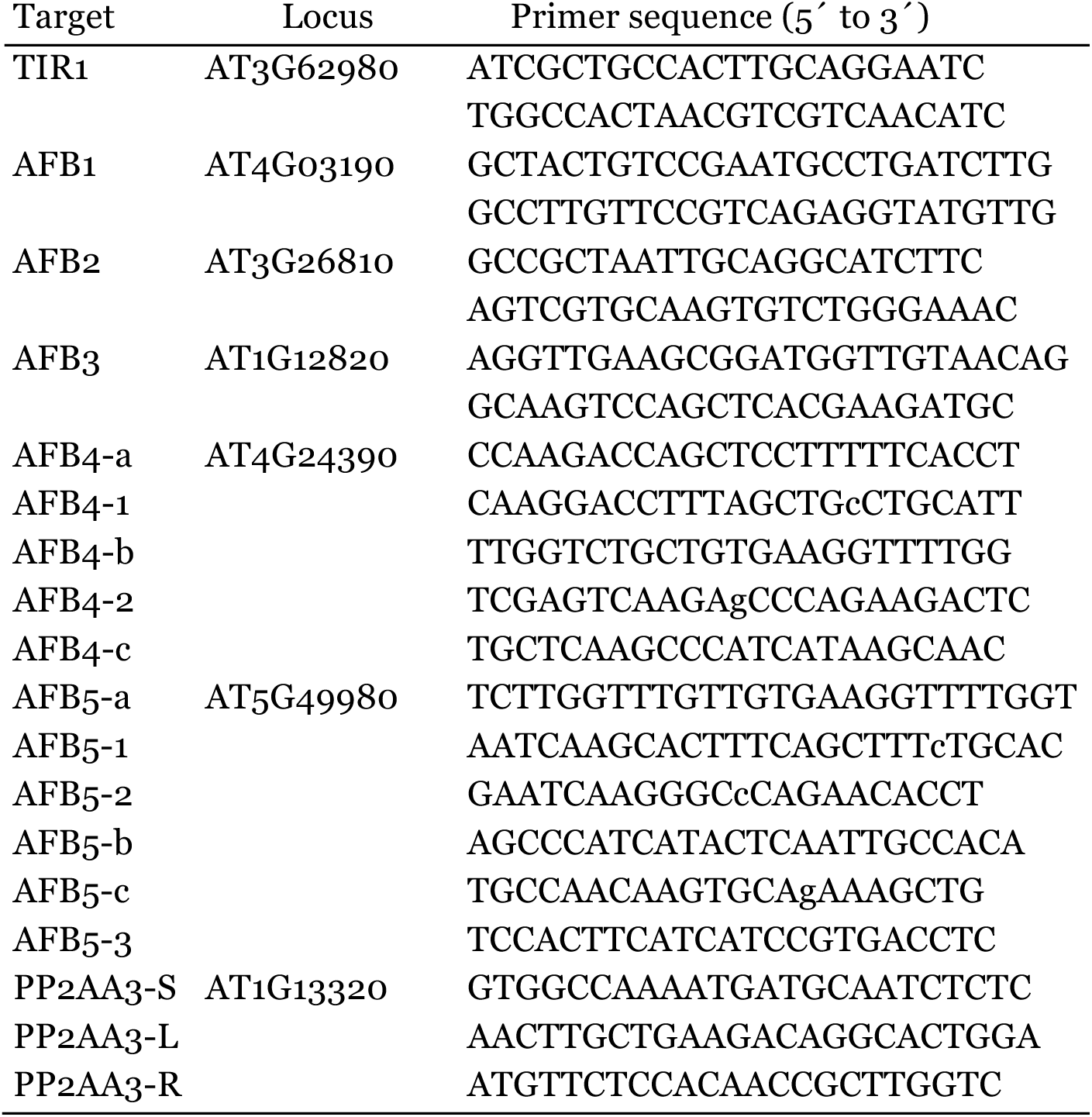

### Results and Discussion

A phylogenetic analysis revealed that the AFB4/AFB5 clade diverged from the TIR1/AFB1-3 clade ~300-400 million years ago whereas the AFB2/AFB3 clade diverged from TIR1/AFB1 ~200 million years ago (Parry *et al*. 2009). Genetic and biochemical studies have demonstrated that members of the TIR1 and AFB2 clades both regulate auxin response but differ in their relative contributions to seedling development (Parry *et al*. 2009). However, the phylogenetically distinct AFB4 group comprised of AFB4 (At4g24390) and AFB5 (At5g49980) in Arabidopsis, has not been characterized in as much detail. Since the corresponding genes have been retained in nearly every seed plant genome sequenced to date, it is likely that they have evolved distinct functions. To explore this possibility we performed a series of experiments focusing on the role of AFB4 and AFB5 during seedling development.

### The AFB4 and AFB5 proteins are auxin receptors

Our first objective was to determine if AFB4 and AFB5 are subunits of SCF complexes. Transgenic lines expressing Myc-tagged versions of AFB4 and AFB5 under the control of the *AFB5* promoter were generated for co-immunoprecipitation experiments. AFB4-Myc and AFB5-Myc were immunoprecipitated from plant extracts with the anti-c-Myc antibody coupled to agarose beads. After washing, the samples were resolved by SDS-PAGE, blotted, and probed with antibodies to the *Arabidopsis* SKP1-related protein ASK1 (Gray *et al*. 1999). A line expressing TIR-Myc was included for comparison (Gray *et al*. 1999). Consistent with their similarity to the TIR1 and AFB1-3 proteins both AFB4 and 5 interact with ASK1 and presumably form an SCF complex (Figure 1A).

To determine whether AFB4 and AFB5 also exhibit the characteristics of auxin receptors, we performed pull-down experiments with the Aux/IAA protein IAA3. Equivalent amounts of total protein extract from AFB4-Myc and AFB5-Myc plants were incubated with GST-IAA3 bound beads in the presence or absence of 50μM IAA. Both AFB4 and AFB5 interact with IAA3 in an auxin dependent manner demonstrating that these proteins function as auxin receptors (Figure 1B).

### AFB4 and AFB5 are the major targets of the picolinate class of auxinic herbicides

The synthetic auxin picloram (4-amino-3,5,6-trichloropicolinic acid) has been well studied for its auxinic herbicidal properties on a variety of plant species (Hamaker *et al*. 1963; Scott and Morris 1970; Chang and Foy 1983). To identify genes required for herbicide response, Walsh and colleagues screened EMS-mutagenized *Arabidopsis* seedlings to identify mutants that were specifically resistant to picolinate auxins (Walsh *et al*. 2006). One of the genes identified in this screen was *AFB5*. Further characterization revealed that the *afb5* mutants were highly resistant to picloram but sensitive to 2,4-D (2,4-dichlorophenoxyacetic acid), a synthetic auxin from the aryloxyacetate class (Walsh *et al*. 2006). In addition, we recently showed that AFB5-Aux/IAA co-receptors selectively bind picloram (Calderon Villalobos *et al*. 2012). To further explore this specificity, we obtained a T-DNA insertion allele of *AFB5* referred to as *afb5-5*. This allele has an insertion in intron 1 that results in the loss of full-length *AFB5* mRNA (Figure 2B and D). In addition we identified two *afb4* mutants with insertions in exon 2 (*afb4-8*) and exon 1 (*afb4-9*) (Figure 2A). Quantitative RT-PCR analysis shows that the *afb4-8* does not produce transcript downstream of the insertion site while transcripts from *afb4-9* plants do not include the first exon (Figure 2C). Thus both alleles are likely to be null mutants. The root growth response of these mutants to picloram was determined and compared to Col-0 and the *tir1-1 afb2-3* double mutant. Consistent with Walsh *et al*. (Walsh *et al*. 2006), *afb5-5* seedlings were strongly resistant to picloram-mediated root growth inhibition (Figure 3A). The *afb4-8* and *afb4-9* were modestly picloram-resistant while *tir1-1 afb2-3* displayed very slight resistance compared to Col-0. We also tested both double mutant combinations and found that *afb4-8 afb5-5* was slightly more resistant than *afb5-5* alone. In addition we tested the response of the *afb4* and *afb5* mutants to the endogenous auxin IAA (Figure 3B). In contrast to the *tir1-1 afb2-3* mutant, *afb4-8, afb5-5*, and the double mutant did not display significant resistance to IAA.

**Figure 3.**
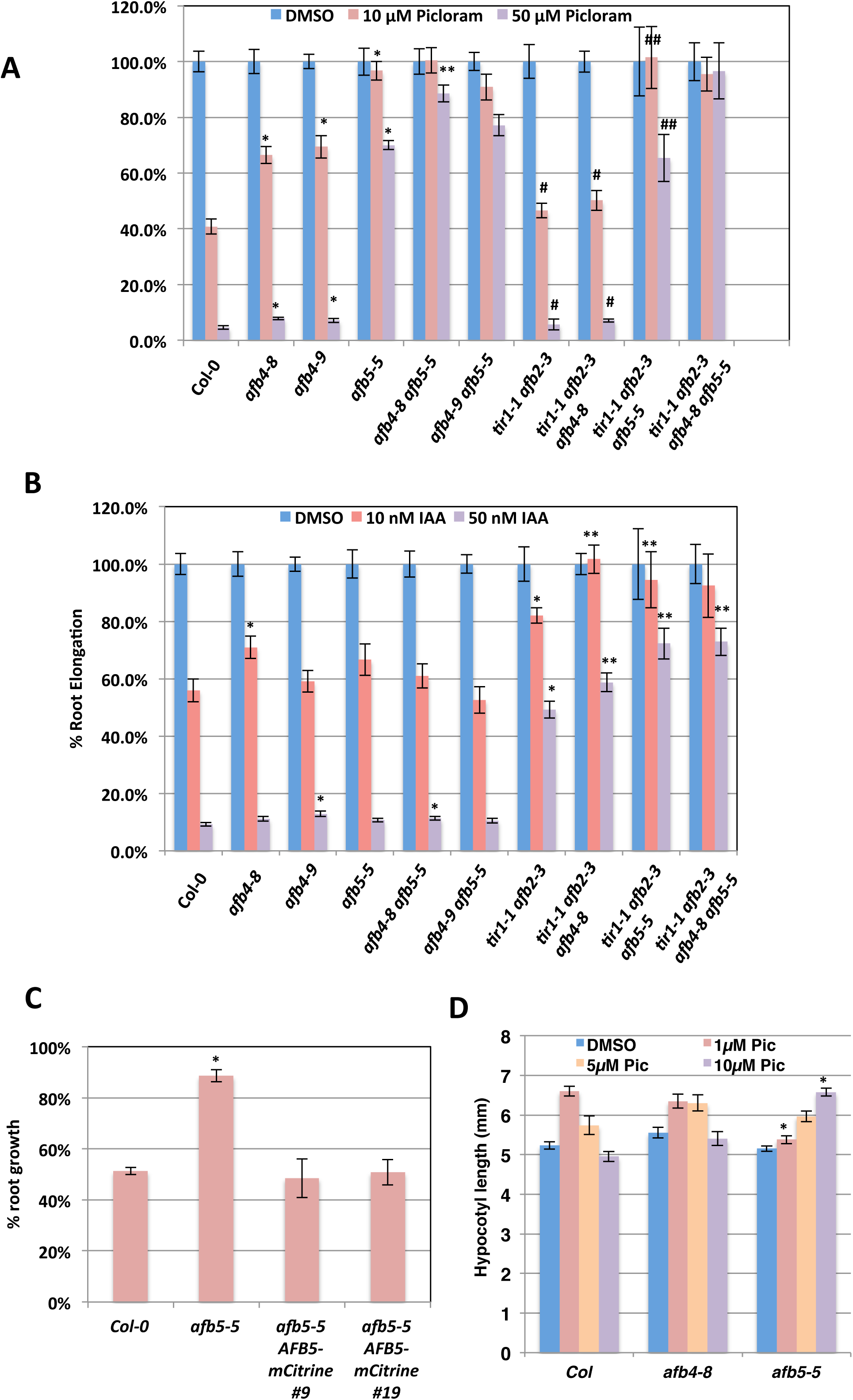
The *afb4* and *afb5* mutants are preferentially resistant to picloram. Five-day old WT and mutant seedlings were transferred to media containing either picloram (A) or IAA (B) and grown another 4 days. Growth is presented as the percent of the DMSO control treatment for each genotype. Error bars represent standard error. (A) *p < 0.05 versus Col-0, **p < 0.05 versus *afb5-5*. # does not exhibit a significant difference with Col-0, ## does not exhibit a significant difference with *afb5-5*. (B) *p <0.05 versus Col-0, ** p<0.05 verus *tir1-1 afb2-3* Student’s *t*-test. (C) Five-day old seedlings for Col-0, *afb5-5*, and two *afb5-5 P_AFB5_:AFB5-mCitrine* lines (T_2_ generation) were transferred to media with or without 10*μ*M picloram and grown for 4 more days before measuring. The *P_AFB5_:AFB5-mCitrine* seedlings were then tested for sensitivity to basta herbicide; measurements from sensitive seedlings were excluded. Results are presented as the percent of the DMSO control treatment for each genotype. *n*= 12, 12, 5, and 5 for Col-0, *afb5-5*, line 9, and line 19, respectively. Error bars represent standard error. * p< 0.05 versus Col-0 and both transgenic lines. (D) Four-day old SD-grown seedlings were transferred to media containing 1*μ*M, 5*μ*M, or 10*μ*M picloram or the equivalent amount of DMSO and grown for two more days before measuring hypocotyl lengths. Error bars represent standard error. *p<0.05 versus Col-0 at the same concentration.

Since we only have a single mutant allele of *AFB5* we sought to confirm that the picloram resistance exhibited by the *afb5-5* allele is due to loss of AFB5. To do this we generated a *P_AFB5_:AFB5-mCitrine* construct and introduced into *afb5-5* plants. Two lines expressing AFB5-mCitrine were chosen for further analysis. The experiment shown in Figure 3C shows that both lines exhibited a wild-type level of response to picloram indicating that loss of AFB5 causes the phenotype.

We also examined the effect of picloram on hypocotyl elongation. Seedlings were grown for 4 days under short day (SD) photoperiods before being transferred to fresh plates containing various concentrations of picloram. As expected based on previous studies, picloram stimulates elongation of Col-0 hypocotyls (Figure 3D)(Chapman *et al*. 2012). In contrast, both *afb4-8* and *afb5-5* are resistant to picloram with *afb5-5* displaying a higher level of resistance. These results demonstrate that the picloram dependent hypocotyl elongation is primarily AFB4/5-dependent.

In a previous study we showed that picloram binds to co-receptor complexes containing AFB5, but not TIR1 (Calderon Villalobos *et al*. 2012). To determine if AFB4 also displays this selectivity, pull-down assays were carried out as before but with the addition of 50μM picloram. Both AFB4 and AFB5 interacted with IAA3 in a picloram-dependent manner whereas the interaction between TIR1 and IAA3 was only slightly affected by picloram (Figure 4A). We also examined the interaction of AFB4 and AFB5 with other auxins in a pulldown experiment (Fig 4B). The results indicate that both proteins also respond to IAA, 2,4-D and 1-NAA. These results suggest a unique specificity of the AFB4 clade for picloram and presumably, related compounds and are consistent with our previous studies showing that the AFB5-IAA7 co-receptor displays selective binding for picloram.

**Figure 4.**
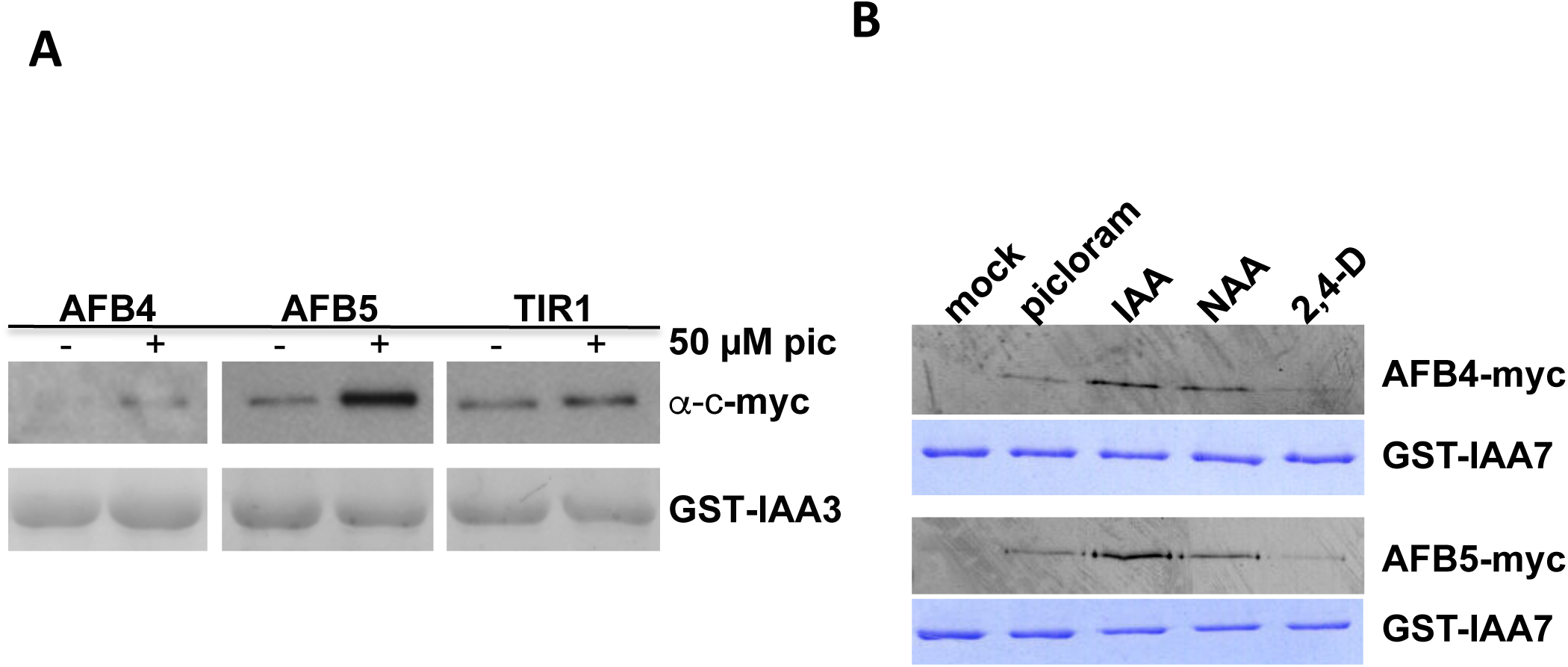
The AFB4 and AFB5 proteins respond to picloram. (A and B) Pull-down reactions were carried out as in figure 1 with 50*μ*M of the indicated auxin. GST-IAA7 loaded was visualized by Coomassie staining.

Taken together, these data indicate that members of the AFB4 clade are the major targets of the picolinate herbicides in Arabidopsis. This finding is particularly important because of the broad use of picloram in agriculture. Identifying the genes that contribute to picloram sensitivity will provide the basis for the development of picloram resistant crops.

### Loss of AFB4 does not result in an obvious seedling phenotype

Previous studies have reported that mutations in *AFB4* confer a pleiotropic phenotype. The *afb4-1* allele was shown to exhibit a variety of growth defects as well as resistance to some pathogens (Hu *et al*. 2012). In another report, the *afb4-2* mutant was reported to have a tall hypocotyl and be auxin hypersensitive (Greenham *et al*. 2011). In contrast *afb4-8* and *afb4-9* do not exhibit any of these qualities. Since these two alleles are nulls, it is clear that AFB4 is not a negative regulator of auxin response. We have shown in subsequent studies that the mutation responsible for the hypocotyl phenotype of *afb4-2* is genetically separable from *AFB4*. In addition, it is our experience that the severe phenotype of the *afb4-1* mutant is unstable suggesting that other factors are contributing to the behavior of this line.

The *afb4-2* mutation does not confer auxin hypersensitivity. However, it is striking that the resulting amino acid substitution, D215N, affects the residue that corresponds to TIR1 D170. In a previous study we showed that the TIR1 D170E mutation does confer auxin hypersensitivity (Yu *et al*. 2013). Because D215N results in loss of a negatively charged residue, whereas D170E does not, we wondered if the *afb4-2* mutation might disrupt AFB4 function. To test this, we performed an *in vitro* pulldown assay with AFB4 and afb4^D215N^ proteins synthesized in a TNT extract. We used IAA7 protein synthesized in *E. coli* for the pulldown. The results shown in Figure 5A,B show that the D215N substitution dramatically reduced recovery of the protein indicating that this mutation does effect function of AFB4.

**Figure 5.**
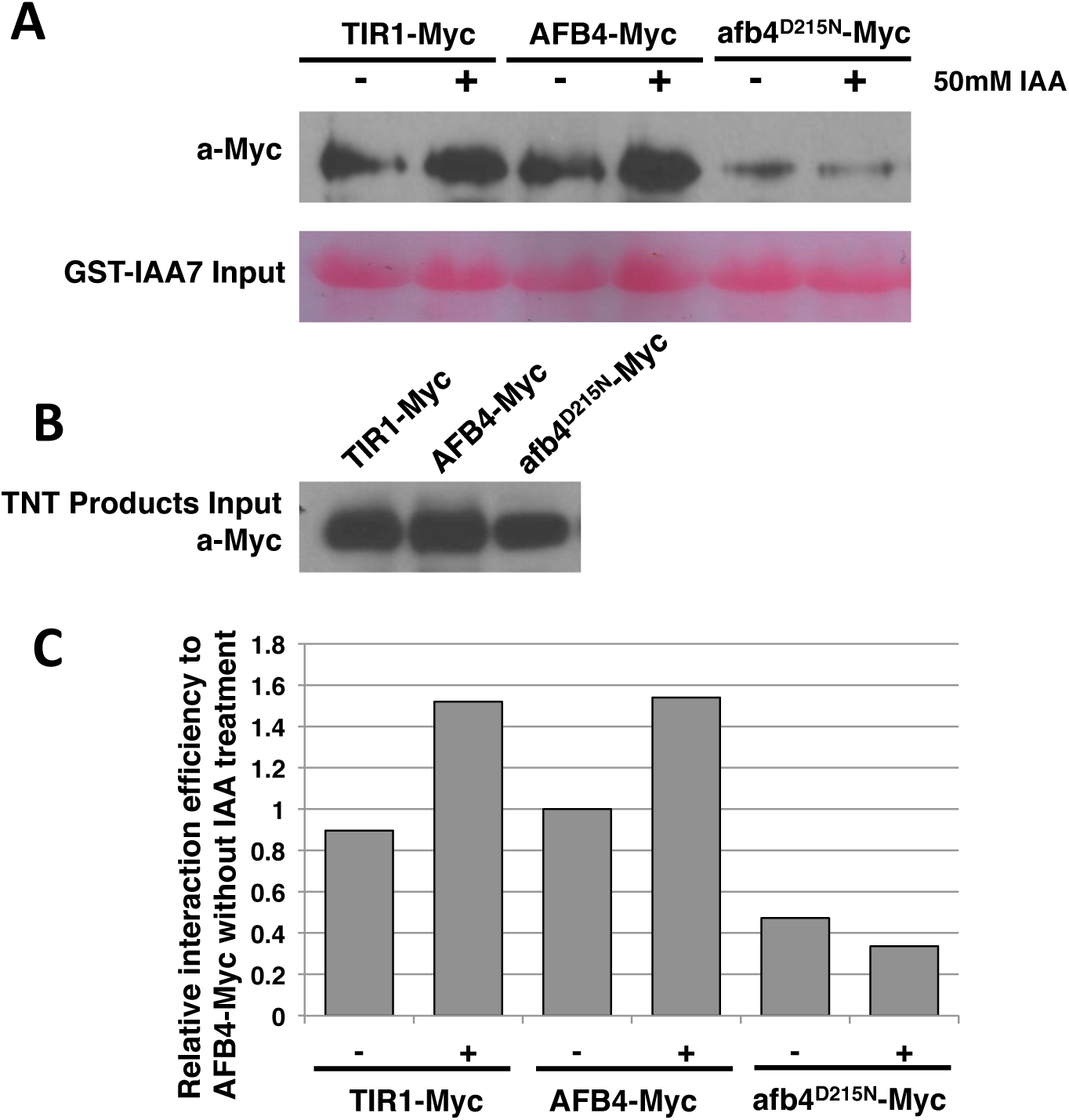
The afb4^D215N^ protein has reduced affinity for IAA7. Equivalent amounts of in vitro translated Myc-tagged TIR1, AFB4, or afb4^D215N^ proteins were incubated with GST-IAA7 protein attached to glutathione-agarose beads in the presence or absence of 50 *μ*M IAA. After washing and elution from the agarose beads, the proteins were separated by polyacrylamide gel electrophoresis and blotted to nitrocellulose membranes. (A) The Myc-tagged receptor proteins were immunodetected using anti-c-Myc antibody and the GST-IAA7 input was visualized by Ponceau S staining. (B) The relative amounts of Myc-tagged proteins added. (C) The quantification of blot band density as presented in (A) by ImageJ. All the values were normalized to AFB4-Myc without IAA treatment.

### Expression of the *AFB4* and *AFB5* genes

To investigate expression of the *AFB4* and *AFB5* genes we measured transcript levels for each of the *T1R1/AFB* genes in tissue collected from 4-day-old seedlings by quantitative RT-PCR. The results in Figure 6 indicate AFB4 and AFB5 are expressed in all tissue types. *AFB4* transcript levels are similar in the root, hypocotyl and cotyledon, whereas the other members of the *T1R1/AFB* family exhibit different levels of expression in cotyledons, hypocotyls and roots. In addition, published transcriptomic data show that *AFB4* is expressed at a relatively low level in most tissues in the plant (Figure 6B) (Schmid *et al*. 2005; Winter *et al*. 2007).

**Figure 6.**
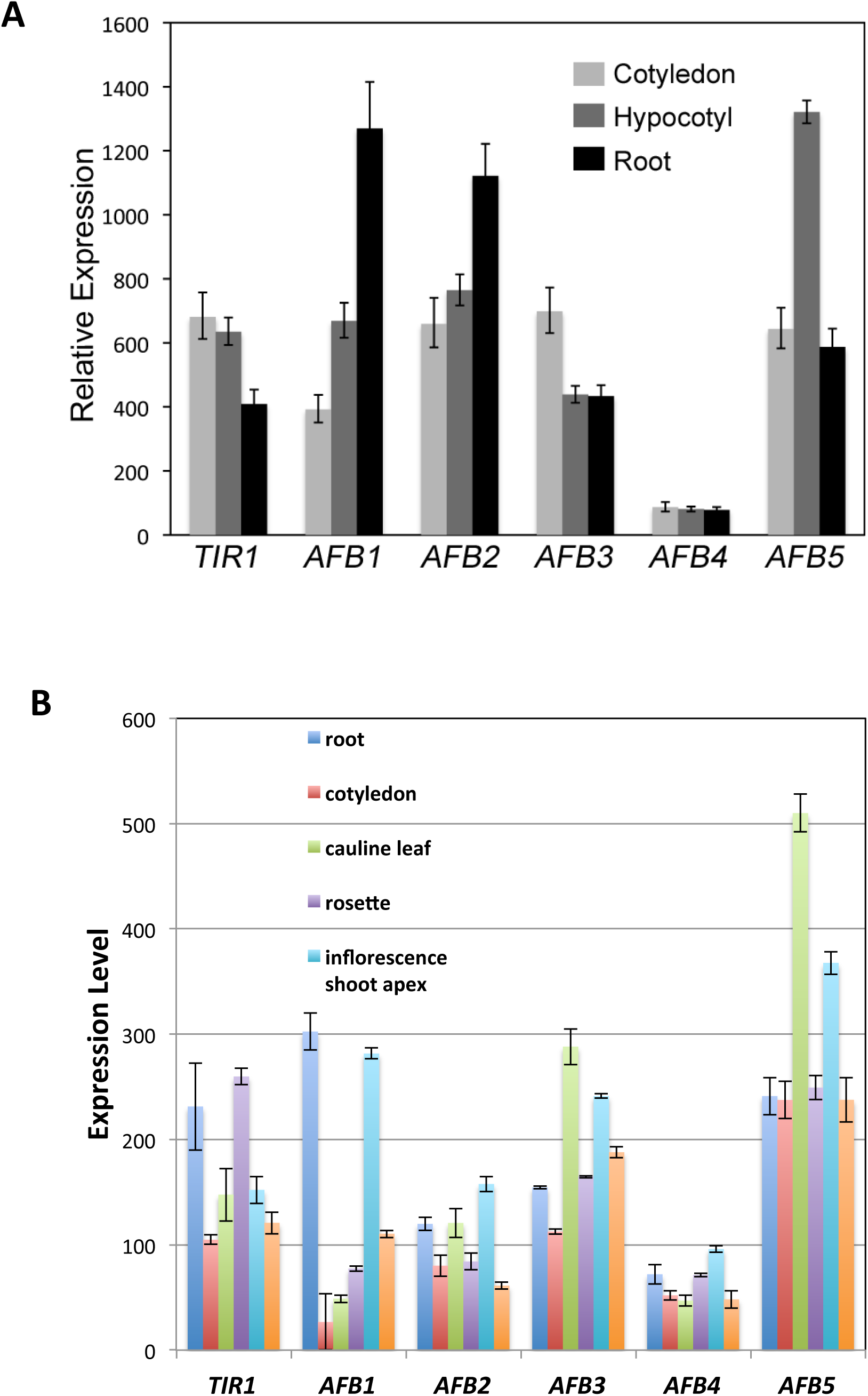
Expression of the *AFB4* and *AFB5* genes. (A) qRT-PCR of *TIR1/AFB* genes in 4-day-old WT seedling tissues grown under short-day (SD) conditions. Primer pairs are listed in Table 1 with AFB4-4 and AFB5-4 being used for those respective genes. Expression is normalized to *PP2AA3* using the PP2AA3-S primer pair. Error bars represent standard error. (B) *TIR1/AFB* expression levels in various tissues. Replotted from (Winter *et al*. 2007).

## Conclusions

In previous studies we demonstrated that TIRI, AFB1, AFB2, AFB3, and AFB5 all bind the Aux/IAA proteins in an auxin dependent manner (Dharmasiri *et al*. 2005; Calderon Villalobos *et al*. 2012). Here we show that AFB4 also functions as an auxin receptor in a manner that is similar to the other members of the family. In addition, we present genetic evidence showing that both AFB4 and AFB5 respond to the synthetic auxin picloram, although the function of AFB5 in picloram response is much greater than that of AFB4. We expect that further genetic studies of the entire family of F-box protein auxin receptors may shed new light on the specialized functions of these proteins.

## Acknowledgments

We thank Nikita Kadakia and Vanessa Peterson for technical assistance. Work in the authors’ lab was supported by a grant from the NIH (GM43644 to ME) and from the NSF (MCB-1122250 to JRE).

